# CNVkit: Copy number detection and visualization for targeted sequencing using off-target reads

**DOI:** 10.1101/010876

**Authors:** Eric Talevich, A. Hunter Shain, Thomas Botton, Boris C. Bastian

## Abstract

Germline copy number variants (CNVs) and somatic copy number alterations (SCNAs) are of significant importance in syndromic conditions and cancer. Massive parallel sequencing is increasingly used to infer copy number information from variations in the read depth in sequencing data. However, this approach has limitations in the case of targeted re-sequencing, which leaves gaps in coverage between the regions chosen for enrichment and introduces biases related to the efficiency of target capture and library preparation.

We present a method for copy number detection, implemented in the software package CNVkit, that uses both the targeted reads and the nonspecifically captured off-target reads to infer copy number evenly across the genome. This combination achieves both exon-level resolution in targeted regions and sufficient resolution in the larger intronic and intergenic regions to identify copy number changes. In particular, we successfully inferred copy number at equivalent to 100-kilobase resolution genome-wide from a platform targeting as few as 293 genes. After normalizing read counts to a pooled reference, we evaluated and corrected for three sources of bias that explain most of the extraneous variability in the sequencing read depth: GC content, target footprint size and spacing, and repetitive sequences. We compared the performance of CNVkit to copy number changes identified by array comparative genomic hybridization. We packaged the components of CNVkit so that it is straightforward to use and provides visualizations, detailed reporting of significant features, and export options for compatibility with other software.

CNVkit is freely availabile from http://github.com/etal/cnvkit.

## 1 Introduction

Copy number changes are a useful diagnostic indicator for many diseases, including cancer. The gold standard for genome-wide copy number is array comparative genomic hybridization (array CGH) (Pinkel et al. 1998; Pinkel and Albertson 2005). More recently, methods have been developed to obtain copy number information from whole-genome sequencing data (Yoon et al. (2009); reviewed by Zhao et al. (2013)). For clinical use, sequencing of genome partitions, such as the exome or a set of disease-relevant genes, is often preferred to enrich for regions of interest and sequence them at higher coverage to increase the sensitivity for calling variants (Dahl et al. 2007). Tools have been developed for copy number analysis of these datasets, as well, including CNVer (Medvedev et al. 2010), ExomeCNV (Sathirapongsasuti et al. 2011), XHMM (Fromer et al. 2012), ngCGH (Gartner et al. 2012), CoNIFER (Krumm et al. 2012), CONTRA (Li et al. 2012), VarScan 2 (Koboldt et al. 2012), EXCAVATOR (Magi et al. 2013) and recent versions of Control-FREEC (Boeva et al. 2011). However, these approaches do not use the sequencing reads from intergenic and, usually, intronic regions, limiting their potential to infer copy number across the genome.

During the target enrichment, targeted regions are captured by hybridization; however, a significant quantity of off-target DNA remains in the library, and this DNA is sequenced and represents a considerable portion of the reads. Thus, off-target reads provide a very low-coverage sequencing of the whole genome, in addition to the high-coverage sequencing obtained in targeted regions. While the off-target reads do not provide enough coverage to call single-nucleotide variants (SNVs) and other small variants, they can provide useful information on copy number at a larger scale.

We developed a computational method for analysis of copy number variants and alterations in targeted DNA sequencing data that we packaged into a software toolkit. This toolkit, called CNVkit, implements a pipeline for CNV detection that takes advantage of off-target sequencing reads and applies a series of corrections to improve accuracy in copy number calling. First, we show that off-target reads map well to the genome and provide a meaningful indicator of copy number. We compare binned read counts in targeted and off-target regions and find that they provide comparable estimates of copy number, albeit at different resolutions. We evaluate several bias correction algorithms to reduce the variance among binned read counts unlikely to be driven by true copy number changes. Finally, we compare copy ratio estimates by the CNVkit method to those of array CGH. In summary, we demonstrate that off-target reads provide reliable copy ratio estimates and their inclusion in analyses maximizes the copy number information obtained from targeted sequencing.

## 2 Results

We evaluated our method on DNA sequencing data from two sets of samples, referred to here as “TR” and “EX”:

- Targeted sequencing (“TR”) comprises 82 melanoma samples, paired tumor and normal tissue from 41 patients, sequenced with a 293-gene target capture protocol.
- Exome sequencing (“EX”) comprises 20 melanoma samples, paired tumor and normal from 10 patients, sequenced with a whole-exome capture protocol.

For each panel of targets, on-target and off-target genomic regions are each partitioned into bins (Table 1) in which unique reads are counted in the initial step of copy number estimation. The read counts and percentages in on-target and off-target regions for each of these samples are shown in Supplemental Table S1. The targets and bin locations used for both sets are included in the CNVKit distribution, along with a test suite which can be executed to repeat our copy number analysis.

**Table 1:** Binning statistics. EX covers a slightly smaller total genomic footprint than TR because most introns are smaller than the minimum size allowed for off-target bins, and thus discarded from the EX bins, while the TR off-target bins span both the introns and exons of non-targeted genes.

| Statistic | TR on-target | TR off-target | EX on-target | EX off-target |
| --- | --- | --- | --- | --- |
| Number of bins | 8216 | 19453 | 301258 | 55064 |
| Total bin footprint | 1,799,531 | 2,848,623,246 | 70,666,905 | 2,479,689,994 |
| Mean bin size | 219.0 | 146,400 | 234.6 | 45,030 |
| Min. bin size | 37 | 10,010 | 115 | 6,000 |
| 1st quartile bin size | 184 | 148,200 | 198 | 11,340 |
| Median bin size | 204 | 149,800 | 228 | 45,030 |
| 3rd quartile bin size | 260 | 151,100 | 269 | 86,870 |
| Max. bin size | 398 | 223,800 | 400 | 135,000 |

### 2.1 On– and off-target read depths similarly reflect copy number

Empirically, we have observed extreme amplifications in off-target areas as sharply demarcated regions with increased read depth (Figure 1A). Since read depth and copy number have been previously shown to be closely correlated (Alkan et al. 2009; Chiang et al. 2009; Xie and Tammi 2009; Yoon et al. 2009), we therefore hypothesized a proportional relationship between read counts and copy number in on-target and off-target bins. The overall agreement in target and off-target bin read counts can be visually verified within selected regions of an individual sample by plotting both values together (Figure 1A).

**Figure 1:**
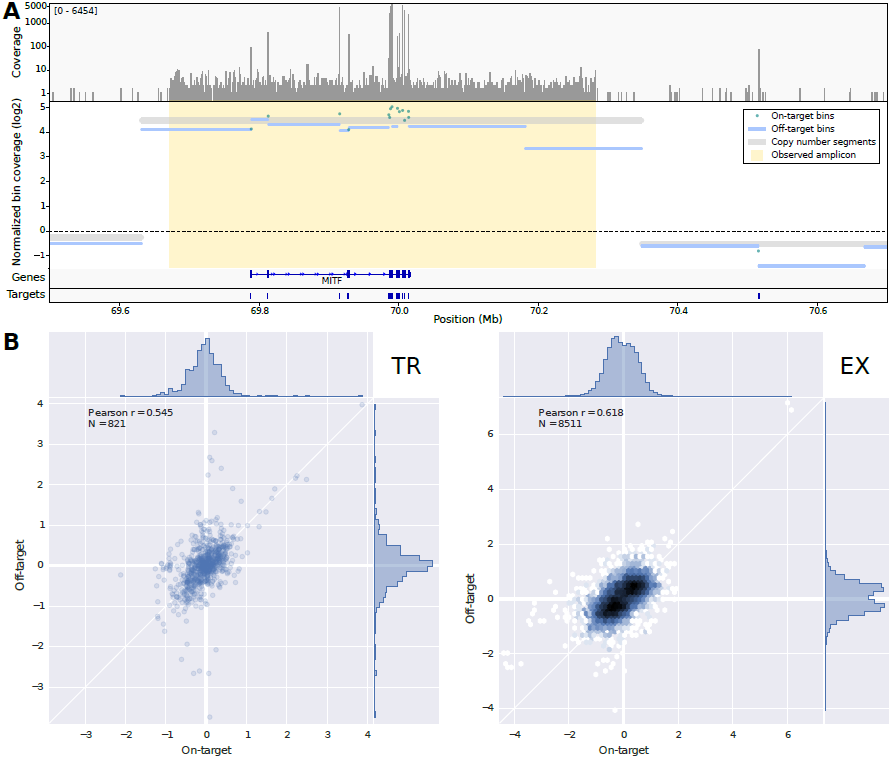
Copy number changes affect read counts in both targeted and off-target intervals. (A) An amplicon including the targeted gene MITF shows similarly increased read depth in both the targeted exons and the adjacent off-target regions. Top row: Coverage depth in and near the MITF region in a melanoma sample, visualized in logarithmic scale in the Integrative Genomics Viewer. Middle: Read counts in on– and off-target bins, log_2_-transformed and median-centered separately. Bottom: RefGene exonic structure of the MITF gene, and baited intervals used for target enrichment. (B) Correlation of mean on– and off-target bin coverages within genes across all samples in the TR and EX sets. Density of data points along each axis is shown as a histogram at the top and right edge of each plot, and as color saturation in the EX plot.

We quantified the level of agreement between on– and off-target read counts more objectively using the TR and EX cohorts. In each cohort we identified the genes containing or adjacent to at least three on– and off-target bins each. For each sample, we performed copy number segmentation with CNVkit and identified the subset of genes in which segmentation indicated a copy number change of at least 0.4-fold at any point within the gene. For each of these genes in this subset, we then calculated the mean of the median-centered log_2_ read counts of the on– and off-target bins separately for each sample. We compared these values from all qualifying genes of all samples and confirmed that the mean on– and off-target log_2_ read counts correlated strongly within genes and appeared to be linearly related across a considerable range (Figure 1B). Thus, off-target read depth provides similar information on copy number status as the on-target read depth.

A substantial amount of noise remains in the relationship between on– and off-target read counts, however. To reduce systematic noise from the copy number signal derived from targeted sequencing data, we next sought to identify and remove extraneous sources of variation in read depth.

#### 2.1.1 Off-target reads reliably map to the genome

In using off-target read counts to estimate copy number, we raised the question of whether the discrepencies in read counts between on– and off-target regions are partly due to unreliable mapping of off-target reads; in particular, whether the mapping quality of off-target reads is significantly less than that of on-target reads.

The mapping quality score (MAPQ) assigned to each aligned read by BWA and similar software indicates the reliability of the read’s position, taking into account read base qualities as well as the alignment scores of the best alignment and secondary or suboptimal alignments, and assigning lower scores for reads mapped to repetitive sequence regions of the reference genome (Li et al. 2008; Li and Durbin 2010). The maximum reported quality score (MAPQ = 60) indicates unambiguous mapping of a read.

For each of the sample cohorts described above (TR, EX), we extracted the mapping qualities of reads in target and off-target regions and compared them to address this question. In the TR samples 98.5% of on-target reads but slightly fewer of the off-target reads (88.3%) were mapped with the maximum quality, while in the EX set there was no overall difference in mapping qualities between on– and off-target reads, with 99.0% mapped with maximum quality in both cases (Figure 2).

**Figure 2:**
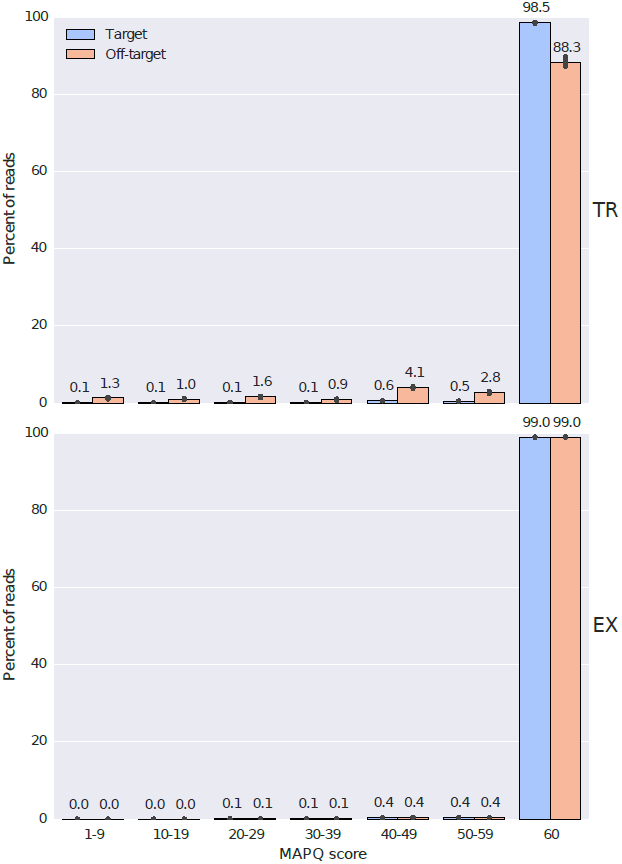
Mapping quality scores of on-target and off-target reads are comparable. For each sample, reads are counted and grouped by MAPQ score within each range labeled on the x-axis, then normalized to the total number of reads obtained from the sample. Bar height indicates the mean of these percentages within each bin; error lines indicate 95% confidence intervals. Scores are shown separately for the targeted (TR) and exome (EX) samples.

### 2.2 Correction of systematic biases affecting read counts

Read depth alone is an insufficient proxy for copy number because of systematic biases in coverage introduced during library preparation and sequencing. For example, others have observed that read depth is affected by GC content, sequence complexity and the sizes of individual targeted intervals (Boeva et al. 2011; Magi et al. 2013; Boeva et al. 2014). Here we describe methods to remove these biases in coverage, and demonstrate the utility of these corrections to improve copy number estimates from targeted sequencing data.

As a first step in correcting the initial copy number estimates obtained from read counts in target and off-target bins, CNVkit normalizes read counts in each bin to the expected read count derived from a reference of normal samples that the user supplies. Normalizing copy number to an appropriate reference is expected to remove much of the biases attributable to GC content, regional targeting density, and repetitive sequence as well as systematic variations introduced by library preparation and sequencing. However, the extent of each of these systematic biases varies from sample to sample (Figure 3), requiring additional correction measures of the residual biases.

**Figure 3:**
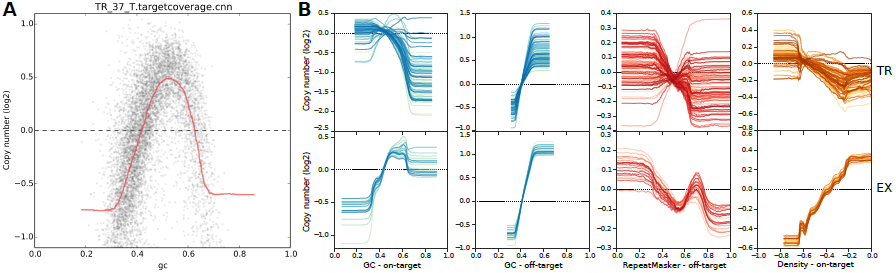
Bin read counts are systematically biased by GC content and other factors. (A) GC coverage bias follows a unimodal distribution in sample TR_37_T. Target bins are sorted according to bin GC content (x-axis), and the uncorrected, median-centered log_2_ bin read counts are plotted (y-axis). A rolling median of the bin log_2_ read counts in order of GC value is drawn in red, showing a systematic deviation from 0 in the selected sample. (B) Trendlines summarize each bias type in each sample. TR and EX samples are shown in the top and bottom rows, respectively. Columns show biases due to GC content in target bins and off-target bins, repeat content in off-target bins, and density bias in target bins.

To account for each of these potentially remaining biases, CNVkit uses a rolling median technique to recenter each on– or off-target bin with other bins of similar GC content, repetitiveness, target size or distance from other targets, independently of genomic location. In the next sections we describe how these biases are addressed in CNVkit.

#### 2.2.1 Read depths vary as a function of regional GC content, targeting density and sequence repeats

DNA regions with higher GC content are less accessible to hybridization and amenable to amplification during library preparation (Aird et al. 2011; Benjamini and Speed 2012). The degree of GC bias may vary between samples due to differences in the quality of each sample’s DNA or efficiency of hybridization between runs. To remove this bias, CNVkit applies the rolling median correction to GC values on both the target and off-target bins, independently.

Repetitive sequences in the genome can complicate read-depth calculations by causing a given read to map similarly well to multiple locations (Magi et al. 2013). Consequently, there is a potential association between sequence repetitiveness and read counts in a region. The strength of this association may vary between samples due to differences in the relative amount of Cot-1 DNA used to block repetitive DNA during library preparation. The human genome sequence provided by the UCSC Genome Bioinformatics Site has repetitive regions masked out by RepeatMasker. CNVKit calculates the proportion of each bin that is masked and uses this information for bias correction. The CNVkit implementation applies the RepeatMasker correction to only the off-target bins. The on-target bins are much smaller, and usually are exonic, and therefore generally have no overlap with repeats. For those on-target bins that were identified as containing repeats (~7% in the TR panel), we found them mostly entirely covered by the repeat, leaving very few intermediate points to infer a continuous trend for correction by the rolling median.

We also noted systematic bias in read depth related to the size of the baited interval and the distance between nearby baited intervals. Two distortions to read depth consistently occured at the edges of each targeted interval (see Methods; Figure 7). The “shoulder” of each interval showed reduced read depth due to incomplete sequence match to the bait, creating a negative bias in the observed read depth inside the interval near each edge; this effect was greatest for short intervals. Some off-target capture also occurred in the “flanks” of the baited interval due to the same mechanism. Where targets are closely spaced or adjacent, this flanking read depth may overlap with a neighboring target, creating a positive bias in its observed read depth. We accounted for these two “density” biases, negative and positive, in a single correction (see Methods). While the density bias was significantly reduced by normalizing each sample to a reference, the density bias was found to vary between samples, likely due to differences in the insert sizes of sequence fragments. Since both these values are calculated in proportion to the target size, the density bias correction also accounts for the bias due to interval size that has been described by others (Magi et al. 2013). Our implementation in CNVkit applies the density correction to only the on-target bins. The negative bias is not expected to occur in off-target regions because those regions are not specifically captured by baits, and the positive bias from neighboring targets is avoided by allocating off-target bins around existing targets with a margin of twice the expected insert size.

#### 2.2.2 Corrections improve copy ratio estimates

We evaluated the effect of each of our bias corrections by comparing the deviation of read counts for on– and off-target bins at each processing step to the final segmented copy number data (Figure 4). For each sample in the TR and EX cohorts, we used CNVkit to perform each of the corrections described above and infer copy number segments. We subtracted the log_2_ copy ratio values of the segments from the median-centered log_2_ read counts of each corresponding on– and off-target bin to obtain the deviations of each bin from our final estimate of true log_2_ copy ratio. From these deviation values, we calculated the standardized median absolute deviation (MAD; a robust measure of scale comparable to standard deviation) across all on– and off-target bins in a given sample. We repeated the MAD calculation at each of the subsequent steps of bias correction: (i) the log_2_ ratios after filtering out poor-quality bins and normalizing to the reference, but before bias corrections; (ii) after GC bias correction; and (iii) after the density and repeat corrections (target and off-target respectively).

**Figure 4:**
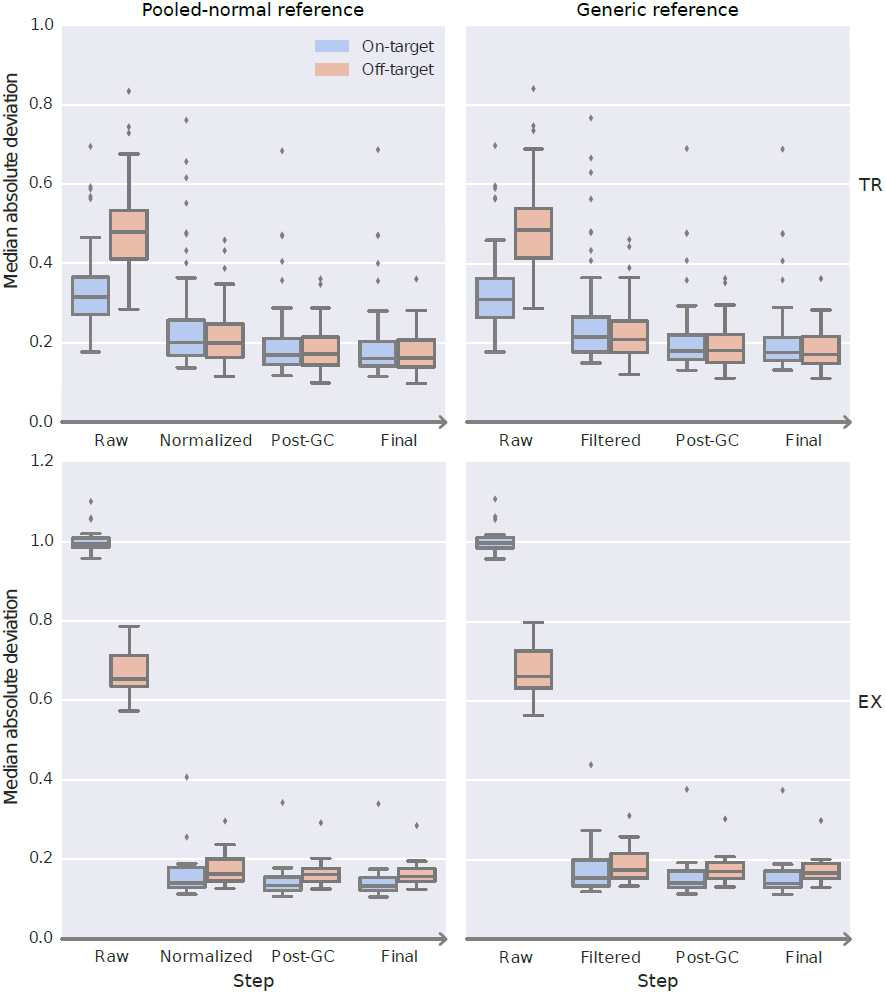
Bias corrections reduce the deviation of bin read depths from the final, segmented copy ratio estimates. Distributions of median absolute deviation of target and off-target bins are shown as boxplots at each step of bias correction for all samples in the TR (top row) and EX (bottom row) sequencing cohorts. Steps shown are the initial median-centered log_2_ read depth (“Raw”), filtering and normalization to a reference (“Normalized”), correction of GC bias (“Post-GC”), and correction of density and repeat biases (“Final”). In the right-side panels, a generic reference was used in place of one constructed from pooled normal samples, and the equivalent second step (“Filtered”) only discards the bins that cover entirely repetitive sequences in the genome.

The first step of normalizing to the reference in CNVkit involves filtering out bins failing certain predefined criteria: those where the reference log_2_ read count is below a threshold (default-5), the spread of read counts among all normal samples in the reference is above a threshold (default 1.0), or the RepeatMasker-covered proportion of the bin is above a threshold (default 99%). To separate the effect of this filtering from that of normalization to a reference of pooled normal samples, we repeated the analysis using a “generic” reference constructed to have a log_2_ read count of 0 assigned to all bins, so that when normalization is performed (but prior to correcting the GC, repeat and density biases) the resulting “copy ratios” are simply equal to the median-centered log_2_ read counts in each bin. Since the generic reference has no meaningful information about read depth derived from normal samples, the effect of using this reference is only to remove the bins with highly repetitive sequence content.

In these results, the MAD values decreased monotonically across all steps, indicating that each step of corrections reduces random deviations from the true copy number signal/value. The spread of MAD values also decreased overall, indicating that the improvements are seen consistently and are reliable; even outlier data points (representing samples with poor overall sequencing quality) were consistently improved. The greatest improvement was seen from filtering repetitive bins, followed by normalization to the pooled reference and GC bias correction. The GC bias correction step reduced the MAD values similarly when using a generic reference or one constructed from normal samples. The improvements from the density and repeat bias corrections were comparatively minor here; however, the MAD values considered are medians aggregated across many bins in a sample, and so this measure would not reflect improvements that are only important in a minority of bins.

We also found that the MAD of the off-target bins was inversely related to the off-target bin size, or equivalently, directly related of the number of reads captured in each off-target bin. Thus, by choosing the off-target bin size to match the average read counts for on-target bins, we ensured that the deviations or random error in the read counts per bin was similar between on– and off-target bins.

### 2.3 Validation by array CGH and FISH assays

The preceding analyses showed that read counts in on– and off-target bins were both consistent indicators of copy number and that consistency can be improved by additional bias corrections. Thus, in CNVkit we combined the bias-corrected log_2_ ratio data from both on– and off-target bins to produce the final estimates of copy number ratio. These copy number ratios were used as input to a segmentation algorithm to infer discrete copy number segments. We used this data to validate the copy ratio calls by CNVkit with respect to two widely used methods for copy number measurement, array CGH and fluorescence in situ hybridization (FISH). For this validation we used the C0902 cell line, derived from a melanoma, which underwent targeted sequencing using a similar protocol as the samples in cohort TR.

CNVkit analysis was performed with default settings and a reference constructed from two unrelated normal tissue samples. We also performed array CGH using a 180K Agilent arraycontaining 174183 probes. Segmentation was performed on the CNVkit bins and raw array CGH probe-level log_2_ ratios using CBS with the same parameters as default in CNVkit.

We compared CNVkit and array CGH copy number ratios across the whole genome (Figure 5A,B). The Pearson correlation coefficient between CNVkit and array CGH copy ratios at all copy number segments overlapping with targeted genes was 96.8% (Figure 5B). The CNVkit and array CGH estimates substantially disagreed at only 1 of the 357 segments. In the gene CEBPA, a 27.5-kilobase loss with a log_2_ copy ratio of -1.5691 is detected by 8 array CGH probes (chr19:33781309-33808811),but the corresponding CNVkit bins showed neutral copy number. These bins cover sequence regions with very high GC content (73–82%) and the CNVkit reference indicated an expected read depth significantly below the genome-wide average, which may have masked any true copy number loss at this locus in the sequencing data.

**Figure 5:**
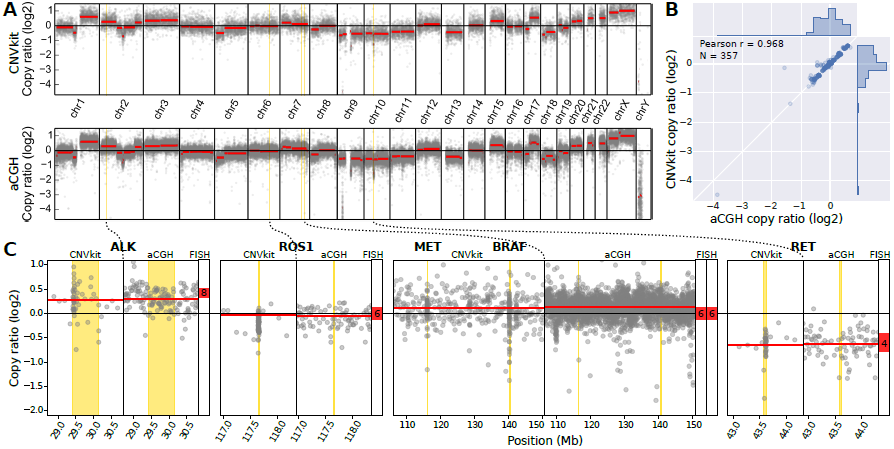
CNVkit copy ratios agree with experimental results array CGH and FISH on cell line DNA. (A) Whole-genome profiles of log_2_ copy ratio by CNVkit (top) and array CGH (bottom) are shown. (B) Segmented copy ratio values over targeted genes correlate closely between array CGH and CNVkit. (C) Genes additionally assayed by FISH are labeled.

Next, we used FISH to determine the absolute copy number at loci harboring cancer-relevant genes: ALK, ROS1, MET, BRAF and RET. We compared the log_2_ ratios obtained by both CNVkit and array CGH to the average signal counts per nucleus obtained by FISH. We transferred the average FISH signal counts into log_2_ copy ratios, by calculating the difference between the log_2_ of their average nuclear signal counts and the log_2_ of the cell’s ploidy, which we determined to be 6n. In all five of the genes assayed by FISH, the copy ratio inferred by CNVkit is close to the true value (Figure 5C).

### 2.4 Software pipeline

We implemented CNVkit as a command-line program,`cnvkit.py,` and reusable Python library,`cnvlib.`

CNVkit uses both the on-target reads and the nonspecifically captured off-target reads to calculate log_2_ copy ratios and segments across the genome for each sample (Figure 6). Off-target bins are calculated from the genomic positions of targets, with the average off-target bin size being much larger than the average on-target bin to match their read counts (Table 1).

**Figure 6:**
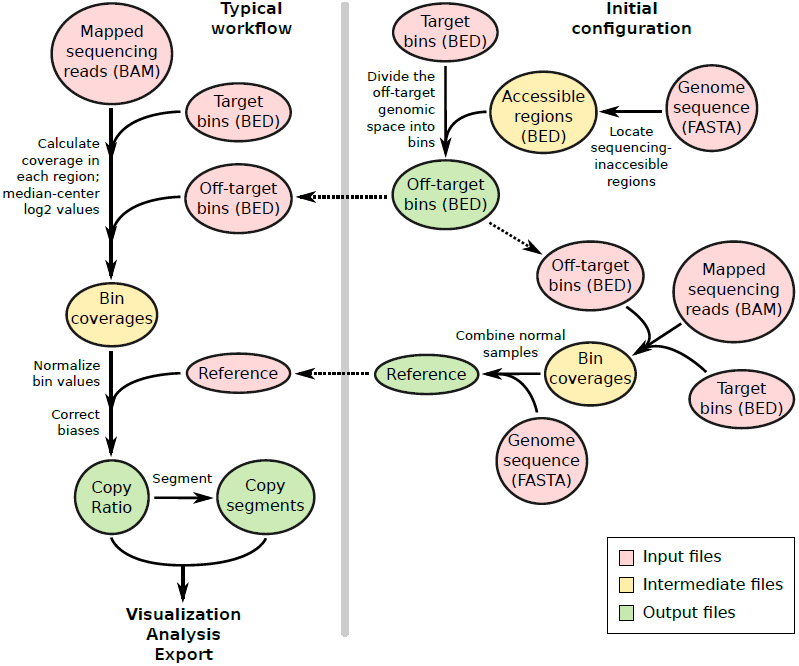
CNVkit workflows. The target and off-target bin BED files and reference file are constructed once for a given platform and can be used to process many samples sequenced on the same platform, as shown in the workflow on the left. Steps to construct the off-target bins are shown at the top-right, and construction of the reference is shown at the lower-right.

Both the on– and off-target locations are then separately used to calculate the mean read depth within each interval. This procedure outputs two separate tabular files of the median-centered log_2_ read depths for on– and off-target bins. The on– and off-target read depths are then combined, normalized to a reference derived from normal samples, corrected for several systematic biases to result in a final table of log_2_ copy ratios. A segmentation algorithm selected by the user is run on the log_2_ ratio values to infer discrete copy number segments. The log_2_ ratios and segments can then be used for visualization and further analyses supported by CNVkit, exported to other formats, and used with third-party software such as the tumor heterogeneity analysis program THetA (Oesper et al. 2013).

These steps are implemented entirely in CNVkit so that the complete workflow can be performed in a reasonable amount of time on a commodity workstation or laptop. The most computationally demanding step, read depth calculation, takes on the order of 30 minutes for an exome or 3 minutes for a 293-gene target panel using a single 3.7GHz CPU. Initial GC region calculation from a genome takes about one minute, and all other steps complete in a few seconds at most. The implementation is designed to be memory-efficient, so that many samples can safely be run in parallel on a single machine.

The input to the program is one or more sequencing read alignments in BAM format (Li et al. 2009) and the bait locations or a pre-built “reference” file. All additional data files used in the workflow, such as GC content and the location of sequence repeats, can be extracted from user-supplied genome sequences in FASTA format using scripts included with the CNVkit distribution. The workflow is not restricted to the human genome, and can be run equally well on other genomes.

## 3 Discussion

We show that off-target reads from partitioned sequencing data are a useful additional indicator of copy number. With an appropriately chosen off-target bin size that matches their read counts to that of on-target bins, we found that the copy ratio estimates in off-target regions can be as reliable or even more reliable than estimates made from on-target regions alone. The combination of on– and off-target reads in copy number estimation can thus achieve both exon-level resolution in targeted regions and greater overall support in the larger intronic and intergenic regions. When samples are sequenced to sufficient depth, target capture design can yield extremely high resolution at specific sites of interest (such as cancer genes and their promoters): Genomic regions around known recurrent structural variation breakpoints, regardless of exon structure, can be tiled to achieve a level of resolution comparable to or even higher than off-the-shelf high-density array CGH.

High-throughput targeted sequencing can be performed with less DNA than is needed for array CGH. Similar to array CGH, samples with low quality or low sequencing coverage (e.g. those with low library complexity, a high number of PCR duplicates and relatively few usable sequencing reads) will have more random variation or “noise” in bin read depths, and overall worse results, due to the fixed size of targeted exons. In these challenging cases it may only be possible to confidently detect CNAs at the chromosome arm level. The off-target bin size used in CNVkit can be increased to improve the fidelity of the copy number signal (at the expense of lower resolution) to identify larger-scale CNAs, particularly those that occur over both on-target and off-target bins.

The copy number reference file plays an important role in the CNVkit pipeline. If the reference is constructed from a large number (e.g. >10) of normal samples, the systematic coverage biases that CNVkit attempts to correct may be largely removed simply by normalization to the reference, and subsequent corrections for GC, repeat and density biases have relatively little effect. However, if using a “generic” reference (with a log_2_ ratio of 0 assigned to all bins), or a reference built from a smaller number of normal samples, or inappropriate normal samples (such as those containing large-scale germline CNVs or sequenced using a different protocol), the additional bias corrections can help compensate for deficiencies in the reference. Ideally, a reference should be constructed specifically for each target capture panel (i.e. set of baits), and match the type of sample (e.g. FFPE-extracted or fresh DNA), and library preparation protocol or kit used. If it is unclear whether a given reference file is suitable for an analysis, one can repeat the analysis with a generic reference and compare the CNAs found with both the original and generic reference for the same samples.

In summary, the CNVkit toolkit provides a suite of programs that maximizes copy number information obtained from targeted DNA sequencing. In addition to making use of both on-target and off-target reads, CNVkit provides robust and efficient implementations of methods to improve estimates of copy number from NGS data.

## 4 Methods

### 4.1 Sequencing

The targeted sequencing cohort (TR) was composed of 41 archived microdissected FFPE melanomas and separately dissected non-lesional “normal” samples from the same patients. All samples were obtained from the archives or the Dermatology Section of the the Departments of Pathology and Dermatology at the University of California San Francisco. The exome sequencing cohort (EX) was composed of 10 fresh frozen melanomas and matching blood samples acquired from Memorial Sloan Kettering Cancer Center. The study was approved by the Institutional Review Boards of both institutions.

Following the manufacturers’ protocols, DNA was extracted using Qiagen DNeasy Blood & Tissue Kit and libraries were prepared for sequencing using the NuGen Ovation Ultralow DR Multiplex System 9-16 (p/n 0331-32). Hybrid capture of the whole exome (EX) and of a targeted panel of 293 cancer-associated genes (TR) was performed using the Agilent SureSelectXT Human All Exon V4+UTRs library (p/n 5190-4638) and the Roche Nimblegen SeqCap EZ Choice Library (p/n 06266339001), respectively. Multiplexed samples were sequenced on an Illumina HiSeq 2500 instrument.

Sequencing reads were aligned to the UCSC reference human genome (hg19; NCBI build 37) with the Burrows-Wheeler Aligner (BWA) version 0.7.5 (Li 2014). PCR duplicates were flagged with Picard MarkDuplicates, and indel realignment and base quality recalibration were performed with the Genome Analysis Toolkit (GATK) to produce the BAM files used as input to CNVkit.

### 4.2 Generating the off-target interval BED files

Genomic intervals for counting off-target reads were initially calculated from the genomic positions of the targeted intervals. The CNVkit antitarget command accepts a list of targeted regions, in Browser Extensible Data (BED) or interval format, and divides the off-target regions between each target into large bins (of 150 kilobases for the TR set and 75kb for EX). The appropriate off-target bin size can be computed as the product of the average target region size and the fold-enrichment of sequencing reads in target regions, such that roughly the same number of reads are mapped to target and off-target bins on average. The preliminary coverage information can be obtained with the script CalculateHsMetrics in the Picard suite (http://picard.sourceforge.net/), or from the console output of the CNVkit `coverage` command when run on the target regions.

Another file listing the sequencing-accessible chromosomal regions can also be provided in order to exclude telomeres, centromeres and other sequencing-inaccessible regions from the off-target intervals when creating the off-target bins. This “access” file has been pre-calculated for the UCSC hg19 genome and is included in the CNVkit distribution; an equivalent file can be generated for any other genome using the bundled script `genome2access.py`, which scans a given FASTA sequence file for long spans of ‘N’ characters in each contig and outputs the coordinates of the regions between them. (Short spans of ‘N’ characters are permitted up to a specified threshold.)

The `antitarget` command divides each contiguous off-target region into equal-sized bins such that the average bin size is as close as possible to the specified bin size. A lower limit on bin size can also be specified to avoid evaluating very small off-target regions, such as some introns, where it is expected that too few reads would be captured to give a reliable estimate of copy number. Once a satisfactory set of off-target bins have been generated and saved as a BED file, the same BED file can be reused with CNVkit for copy number analysis of other samples prepared with the same library preparation protocol and sequenced on the same platform.

### 4.3 Estimation of copy number by read depth

The log_2_ mean read depth of each on– or off-target bin is computed for a given sample using the `coverage` command. Given an alignment of reads to the reference genome, in BAM format, and a list of the on– or off-target bins, in BED or Picard interval format, for each bin the read depths at each base pair in the bin are calculated and summed using pysam, a Python interface to samtools (Li et al. 2009), and then divided by the size of the bin. The output is a table of the average read depths in each of the given bins, log_2_-transformed and centered to the median read depth of all autosomes.

To produce the input BAM file, we recommend that an aligner such as BWA-MEM be used with the option to mark secondary mappings of reads, and that PCR duplicates be flagged with a program such as SAMBLASTER (Faust and Hall 2014) or Picard MarkDuplicates, so that CNVkit will skip these reads when calculating read depth.

### 4.4 Construction of a pooled reference from a panel of normal samples

The `reference` command combines the bin read depths of multiple, ideally normal-tissue samples into a single copy-number reference, which can then be used to correct other individual samples. The output reference dataset contains, for each bin, a weighted average of the log_2_ read depths from the input samples, calculated as Tukey’s biweight location, and the spread or statistical dispersion of read depth values, calculated as Tukey’s biweight midvariance (Lax 1985; Randal 2008).

If a FASTA file of the reference genome is given, the GC content of the corresponding genomic region, calculated as the proportion of “G” and “C” characters among all “A”, “T”, “G” and “C” characters in the subsequence, ignoring “N” and any other ambiguous characters. For efficiency, the samtools FASTA index file (.fai) is used to locate the binned sequence regions in the FASTA file.

For the human genome and many model organism genomes, repetitive sequences have been identified using RepeatMasker and masked out with lowercase characters in the genome sequences provided for download by the UCSC Genome Bioinformatics Site (http://genome.ucsc.edu/) and the RepeatMasker website (http://repeatmasker.org). CNVkit calculates the fraction of each bin that is masked and records this fraction in an additional column in the reference file. When using the reference to normalize and correct individual samples, CNVkit can then filter out bins covering highly repetitive genomic regions, or those with very low average coverage or high spread in the reference.

As with the target and off-target BED files, once a satisfactory reference file has been generated, it can be reused with CNVkit for copy number analysis of other samples sequenced with the same platform and protocol.

### 4.5 Calculation of “density” bias

Two consistent biases in read depth occur related to the size and regional density of baited intervals, which we refer to here as the “density” biases: a negative bias at the interval “shoulder” and a positive bias in the interval “flank” regions. These biases occur within a distance of the interval edges equal to the sequence fragment size. For simplicity (since only a monotonic function of the actual read depth bias is needed for the bias correction to work properly, rather than the magnitude of the bias itself), the effect of the biases is modeled as a linear decrease in read depth from inside the target region to the same distance outside (Figure 7).

**Figure 7:**
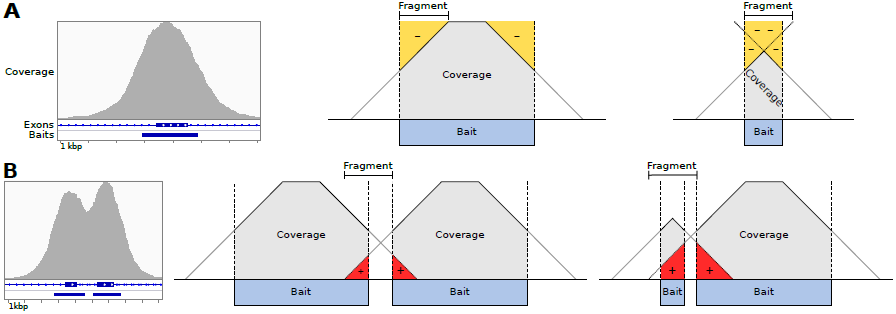
Baited region size and spacing affect read depth systematically. (A) Example of typical coverage observed at a targeted exon, as viewed in IGV, and simplified geometric models of the negative coverage biases (yellow) that can occur as a function of the relative sizes of sequence fragments and the baited region. (B) Coverage observed at two neighboring targeted exons, and models of the positive coverage biases (red) that can occur where intervals are separated by less than half the insert size of sequence fragments.

Letting *i* be the average insert size and *t* be the target interval size, the negative bias at interval shoulders is calculated as *i/*4*t* at each side of the interval, or *i/*2*t* for the interval (Figure 7A). When the interval is smaller than the sequence fragment size, the portion of the fragment extending beyond the opposite edge of the interval should not be counted in this calculation. Thus, if *t <i*, the negative bias value must be increased (absolute value reduced) by 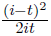.

Additionally letting *g* be the size of the gap between consecutive intervals, the positive bias that occurs when the the gap is smaller than the insert size (*g < i*) is 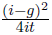(Figure 7B). If the target interval and gap together are smaller than the insert size, the reads flanking the neighboring interval may extend beyond the target, and this flanking portion beyond the target should not be counted. Thus, if *t* + *g* < *i*, the positive value must be reduced by 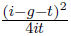.

These values are combined into a single value by subtracting the estimated shoulder bias from the flank bias. The result is a negative number between -1 and 0, or 0 for target with immediately adjacent targets on both sides. Thus, dividing a large targeted interval into a consecutive series of smaller targets does not change the net “density” calculation value.

### 4.6 Correction of coverage biases

The `fix` command combines target and off-target copy number tables for a single sample, removes bins failing predefined criteria, subtracts the reference log_2_ read depth from each bin to yield the log_2_ ratio of observed to expected copy number, corrects for systematic biases in bin coverage (see below), and finally re-centers the corrected copy ratios.

Bins are removed according to hard thresholds applied to the additional fields in the reference file: when the reference coverage is too low, the spread of read depth values in the normal panel is too high, or repeat-masked fraction of the bin is too high.

For each of the biases (GC, density, repeat-masked fraction), the bias value is calculated for each bin. Bins are sorted by bias value. A rolling median is then calculated across the bin log_2_ ratios ordered by bias value to obtain a midpoint log_2_ ratio value representing the expected bias for each bin. This value is then subtracted from the original bin log_2_ ratio for the given sample to offset the observed bias. Local regression (LOWESS) (Cleveland 1979) and a Kaiser window function (Kaiser and Schafer 1980) were also evaluated in place of the rolling median to estimate the trend due to bias; all three functions produced similar fits on sample data, and rolling median was chosen as the default for its simplicity and robustness.

The GC content and repeat-masked fraction of each bin are calculated during generation of the reference from the user-supplied genome. Density biases are calculated from the start and end positions of a bin and its neighbors within a fixed window around the target’s genomic coordinates, as described in the previous section. In the common case of no other targets within the window surrounding the given target, there is a one-to-one correspondence of the density value to target size. Thus, interval size (the difference between the interval start and end coordinates) is not used as an additional bias correction by default, as it is redundant with the density correction.

In the case of many similarly sized target regions, there is the potential for the bias value to be identical for many targets, including some spatially near each other. To ensure that the calculated biases are independent of genomic position, the probes are randomly shuffled before being stably sorted by bias value.

### 4.7 Segmentation

The sample’s corrected bin-level copy ratio estimates can be segmented into discrete copy-number regions using the `segment` command. The default segmentation algorithm used is cicular binary segmentation (CBS) (Olshen et al. 2004), via the R package PSCBS (Olshen et al. 2011). Alternatively, the HaarSeg algorithm (Ben-Yaacov and Eldar 2008) or Fused Lasso (Tibshirani and Wang 2008) can be used in place of CBS. In either case, the segmentation output is in a tabular format similar to the copy number and copy ratio tables used by CNVkit.

### 4.8 Cell line sequencing, array CGH and FISH

### 4.9 Targeted sequencing

Targeted sequencing on C0902 cells was performed using a panel of targeted genes different from those used in the TR set, but similarly focused to about 300 genes. The target panel covered 6915 baited regions; binning yielded 8699 target bins and 19797 off-target bins, for a total of 28496 bins.

#### 4.9.1 Comparative genomic hybridization

DNA was extracted from the C0902 cell line using a Flexigene DNA extraction kit (Qiagene, Germantown, MD, USA) according to manufacturer’s protocol. Array CGH was carried out with 1000 ng of genomic DNA on Agilent 4x180K microarrays (Agilent, Santa Clara, CA, USA). The raw microarray images were processed with Agilent Feature Extraction software. Probe copy ratio values were then converted to the CNVkit format with a Python script, skipping unassigned contigs and “dummy” probes.

#### 4.9.2 Fluorescence in situ hybridization (FISH)

ROS1 and RET break-apart FISH probes were labeled commercial probes purchased from Kreatech Diagnostics (Amsterdam, The Netherlands). ALK probes were from Abbott Molecular (Des Plaines, Illinois, USA). BRAF, MET and NTRK1 break-apart FISH probes were prepared from BAC clones using standard procedures, and labeled with Fluorolink Cy3-dUTP (GE Healthcare, Waukesha, WI, USA) and ChromaTide Alexa Fluor 488-5-dUTP (Life Technologies, Gaithersburg, MD, USA).

After 10 minutes of incubation at 37°C in a hypotonic solution of cell culture medium/distilled water (5:7), C0902 cells were fixed in methanol/glacial acetic acid (3:1) and dropped on a slide. After two days of aging, the slide was treated with RNAse and proteinase K before the probes were hybridized. The number and localization of the hybridization signals was assessed in interphase nuclei with well-delineated contours using a Zeiss fluorescence microscope.

### 4.10 Visualization

Plots were generated using the Python programming language and Biopython (Cock et al. 2009), matplotlib (http://matplotlib.org), Seaborn (https://github.com/mwaskom/seaborn) and CNVkit software libraries. Figures were arranged and labeled using Inkscape (http://inkscape.org).

## 5 Data access

Data files used for the evaluation of CNVkit are available at http://github.com/etal/cnvkit-examples.

## 6 Acknowledgements

We thank Thomas Botton for support in the analysis of cell line data, Henrik Bengtsson for helpful suggestions on the use of PSCBS, and members of the Bastian lab for comments on the manuscript and feedback during the development of CNVkit.

This work was supported by the National Institutes of Health [R01 CA131524, P01 CA025874 to B.C.B.].

## 8 Supplemental material

Supplemental Table S1: Sequencing metrics

